# How does the presence of neural probes affect extracellular potentials?

**DOI:** 10.1101/318741

**Authors:** Alessio Paolo Buccino, Miroslav Kuchta, Karoline Horgmo Jæger, Torbjørn Vefferstad Ness, Pierre Berthet, Kent-Andre Mardal, Gert Cauwenberghs, Aslak Tveito

## Abstract

**Abstract:** *Objective:* Mechanistic modeling of neurons is an essential component of computational neuroscience that enables scientists to simulate, explain, and explore neural activity. The conventional approach to simulation of extracellular neural recordings first computes transmembrane currents using the cable equation and then sums their contribution to model the extracellular potential. This two-step approach relies on the assumption that the extracellular space is an infinite and homogeneous conductive medium, while measurements are performed using neural probes. The main purpose of this paper is to assess to what extent the presence of the neural probes of varying shape and size impacts the extracellular field and how to correct for them.

*Approach:* We apply a detailed modeling framework allowing explicit representation of the neuron and the probe to study the effect of the probes and thereby estimate the effect of ignoring it. We use meshes with simplified neurons and different types of probe and compare the extracellular action potentials with and without the probe in the extracellular space. We then compare various solutions to account for the probes’ presence and introduce an efficient probe correction method to include the *probe effect* in modeling of extracellular potentials.

*Main results:* Our computations show that microwires hardly influence the extracellular electric field and their effect can therefore be ignored. In contrast, Multi-Electrode Arrays (MEAs) significantly affect the extracellular field by magnifying the recorded potential. While MEAs behave similarly to infinite insulated planes, we find that their effect strongly depends on the neuron-probe alignment and probe orientation.

*Significance:* Ignoring the *probe effect* might be deleterious in some applications, such as neural localization and parameterization of neural models from extracellular recordings. Moreover, the presence of the probe can improve the interpretation of extracellular recordings, by providing a more accurate estimation of the extracellular potential generated by neuronal models.

## 1 Introduction

Huge efforts have been invested in computational modeling of neurophysiology over the last decades. This has led to the development and public distribution of a large array of realistic neuron models, for example from the Blue Brain Project (`bbp.epfl.ch` [1, 2]), the Allen-Brain Institute brain cell database (`celltypes. brain-map.org` [3]), and the Neuromorpho database (`neuromorpho.org` [4, 5]). As experimental data become available, these models become both more elaborate and more accurate. However, some of the assumptions underlying the most commonly used models may not allow the accuracy necessary to obtain good agreements between models and experiments. For instance, it was pointed out in *Tveito et al.* [6] that assumptions underlying the classical cable equation and the associated method for computing the extracellular potential, lead to significant errors both in the membrane potential and the extracellular potential. In the present paper we investigate whether the classical modeling techniques used in computational neurophysiology are sufficiently accurate to reflect measurements obtained by different types of probes, such as microwires/tetrodes, and larger silicon Multi-Electrode Arrays (MEAs). Traditionally, these devices are not represented in the models describing the extracellular field, and our aim is to see if this omission introduces significant errors and how this mismatch could be accounted for in modeling of extracellular activity.

The most widely accepted and used modeling framework for computing the electrophysiology of neurons is the *cable equation* [7, 8, 9, 10, 11], which is used to find current and membrane potentials at different segments of a neuron. One straightforward and computationally convenient way to model the extracellular electric potential generated by neural activity is to sum the individual contributions of the transmembrane currents (computed for each segment) considering them as point current sources or line current sources [7, 11] using volume conductor theory. Although this approach represents the gold standard in computational neuroscience, there are some essential assumptions that need to be discussed. First, *i)* the neuron is represented as a cable of discrete nodes and the continuous nature of its membrane is not preserved. Second, *ii)* when solving the cable equation, the extracellular potential is neglected, but the extracellular potential is computed *a-posteriori.* Third, and foremost, *iii)* when computing extracellular potentials, the tissue in which the neuron lies is modeled as an infinite medium with homogeneous properties. The validity of these assumptions must be addressed in light of the specific application under consideration. The first assumption i) can be justified by increasing the number of nodes in the model, but assumption ii) is harder to relax since it means that the model ignores ephaptic effects. Therefore, this assumption has gained considerable attention [12, 13, 14, 15, 16, 17, 6]. However, the main focus of the present paper is assumption *iii*). More specifically our aim is to study the effect of the physical presence of a neural probe on the extracellular signals. Can it be neglected in the mathematical model, or should it be included as a restriction on the extracellular domain? Specifically, is the conventional modeling framework, ignoring the effect of the probes, sufficient to yield reliable prediction of extracellular potentials? Finally, what can modelers do in order to represent and include the effect of recording probes?

In order to investigate this question, we have used the Extracellular-Membrane-Intracellular (EMI) model [18, 6, 19]. The EMI model allows for explicit representation of both the intracellular space of the neuron, the cell membrane and the extracellular space surrounding the neuron. Therefore, the geometry of neural probes can be represented accurately in the model. We have run finite element simulations of simplified pyramidal cells combined with different types of probes, such as microwires/tetrodes, and larger silicon Multi-Electrode Arrays (MEAs).

Our computations strongly indicate that the effect of the probe depends on several factors; small probes (microwires) have little effect on the extracellular potential, whereas larger devices (such as Multi-Electrode Arrays, MEAs) change the extracellular potential quite dramatically, resembling the effect of a non-conductive infinite plane in the proximity of the neuron. The effect, however, depends on the neuron-probe alignment and orientation. We then compare the EMI results with conventional cable equation-based techniques, such as the current summation approach [11, 19], the hybrid solution [20, 21, 22, 19], and the method of images [23, 24] and introduce the probe correction method, which allows to reach a hybrid solution accuracy leveraging on a pre-mapping of the probe-specific effect and the reciprocity principle.

The results may aid in understanding experimental data recorded with MEAs, it may improve accuracy when extracellular potentials are used to parameterize membrane models as advocated in [25], and to localize and classify neurons from MEA recordings [26, 27].

The rest of the article is organized as follows: in Section 2 we describe the methods used throughout the paper, with particular focus on the EMI model (§2.1), the meshes (§2.2), the finite element framework (§2.3), and modeling approaches used for comparison (§2.4). In Section 3 we present our findings related to the effect of probes of different geometry on the extracellular recordings (§3.1), the variability of our simulations depending on geometrical parameters of the mesh (§3.2), before comparing them with results obtained from other computational approaches (§3.3) and the relative computational costs of these methods (§3.4). Finally, we discuss and contextualize the work in Section 4.

## 2 Methods

In this section we introduce the modeling frameworks used to investigate the effect of the probes on the extracellular potential. In particular we first describe the EMI model, the meshes, and the membrane and finite element modeling. Then, we describe the conventional modeling based on the cable equation solution: the current summation approach (CS), the hybrid solution (HS) and the method of images (MoI). Finally, we introduce the probe correction method (PC), which reaches the hybrid solution accuracy in a more efficient and computationally-cheap way.

### 2.1 The Extracellular-Membrane-Intracellular model

The purpose of the present report is to estimate the effect of introducing a probe in the extracellular domain on the extracellular potential. This can be done using a model discussed in [28, 29, 18, 30, 6] referred to as the EMI-model. In the EMI-model the Extracellular space surrounding the neuron, the Membrane of the neuron and the **I**ntracellular space of the neuron are all explicitly represented in the model. The model takes the form

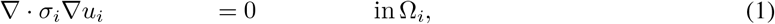

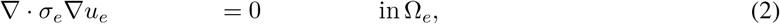

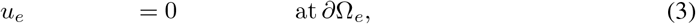

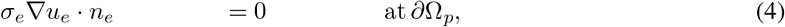

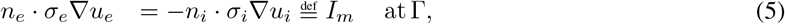

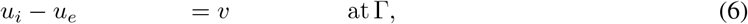

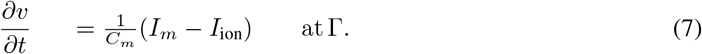

In the simplified geometry sketched in Figure 1, Ω denotes the total computational domain consisting of the extracellular domain Ω_*e*_ and the intracellular domain Ω_*i*_, and the cell membrane is denoted by Γ. *n_i_* and *n_e_* are the vectors normal to Γ pointing to the intra- and extracellular domains, respectively. *u_i_* and *u_e_* denote the intra- and extracellular potentials, and *v* = *u_i_* − *u_e_* denotes the membrane potential defined at the membrane Γ. The intra- and extracellular conductivities are given respectively by *σ_i_* and *σ_e_* and in this work we assume that the quantities are constant scalars. The cell membrane capacitance is given by *C_m_*, and the ion current density is given by *I*_ion_. *I*_m_ is the total current current escaping through the membrane.

**Figure 1:**
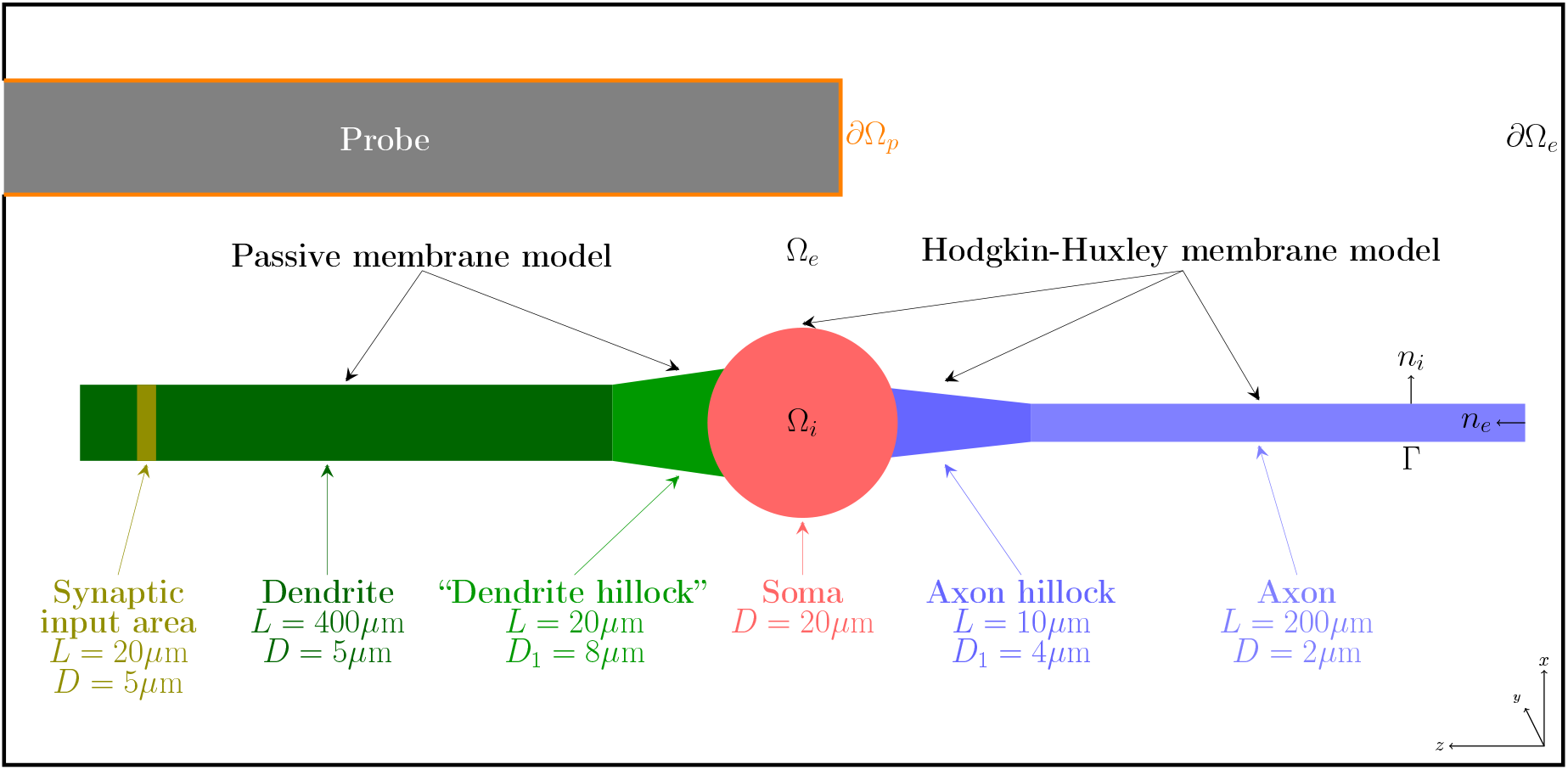
Sketch of the simplified neuron geometry and its surroundings. The intracellular domain is denoted by Ω_*i*_, the cell membrane is denoted by Γ, and the extracellular domain is denoted by Ω_*e*_. The boundary of the probe is denoted by *∂Ω_*p*_* and the remaining boundary of the extracellular domain is denoted by *∂*Ω_*e*_. The normal vector pointing out of Ω_*i*_ is denoted by *n_e_*, and *n_e_* denotes the normal vector pointing out of Ω_*e*_. *L* and *D* are the length and diameter of neural segments, respectively, and *D*_1_ is the diameter of the *hillocks* in correspondence of the soma. In our simulations, we consider three types of probe geometry (see Figure 2). Note that the probe interior is *not* part of the computational domain.

The EMI model is here considered with grounding (Dirichlet) boundary conditions, i.e. *u_e_* = 0, on the boundary of the extracellular domain (*∂*Ω_*e*_) while insulating (Neumann) boundary conditions, i.e. *σ_e_*ᐁ*u_e_* · *n_e_* = 0 were prescribed at the surface of the probe (*∂*Ω_p_). Note that the latter is a suitable boundary condition also for the conducting surfaces of the probe [31, 24]. The resting potential (see Table 1) is used as initial condition for *v*.

### 2.2 Meshes

In order to implement the EMI model described above, the computational domain was discretized by unstructured tetrahedral meshes generated by `gmsh` [32]. We used a simplified neuron model similar to a ball–and–stick model [33, 34], with a spherical soma with 20 μm diameter – whose center is in the origin of the axis – an apical dendrite of length *L*_d_ = 400 μm and diameter *D*_d_ = 5 μm in the positive *z* direction and an axon of length *L*_d_ = 200 μm and diameter *D*_d_ = 2 μm in the negative *z* direction. Both the axon and the dendrites are connected to the soma via a tapering in the geometry. On the dendritic side, the diameter at the soma is 8 μm and it linearly reduces to 5 μm in a 20 μm portion. On the axonal side, the *axon hillock* has a diameter of 4 μm at the soma and it is tapered to 2 μm in 10 μm.

The neuron was placed in a box with and without neural probes to study the effect of the recording device on the simulated signals. We used three different types of probes:

**Microwire**: the first type of probe represents a microwire type of probe (or tetrode). For this kind of probes we used a cylindrical insulated model with 30 μm diameter. The extracellular potential, after the simulations, was estimated as the average of the electric potential measured at the tip of the cylinder. The microwire probe is shown in Figure 2A alongside with the simplified neuron.
**Neuronexus (MEA):** the second type of probe model represents a commercially available silicon MEA (A1x32-Poly3-5mm-25s-177-CM32 probe from Neuronexus Technologies), which has 32 electrodes in three columns (the central column has 12 recording sites and first and third columns have 10) with hexagonal arrangement, a *y*-pitch of 18 μm, and a *z*-pitch of 22 μm. The electrode radius is 7.5 μm. This probe has a thickness of 15 μm and a maximum width of 114 μm, and it is shown in Figure 2B.
**Neuropixels (MEA):** the third type of probe model represents the Neuropixels silicon MEA [35]. The original probe has more than 900 electrodes over a 1 cm shank, it is 70 μm wide and 20 μm thick. In our mesh, shown in Figure 2C we used 24 12×12 μm recording sites arranged in the *chessboard* configuration with an inter-electrode-distance of 25 μm [35].

**Figure 2.**
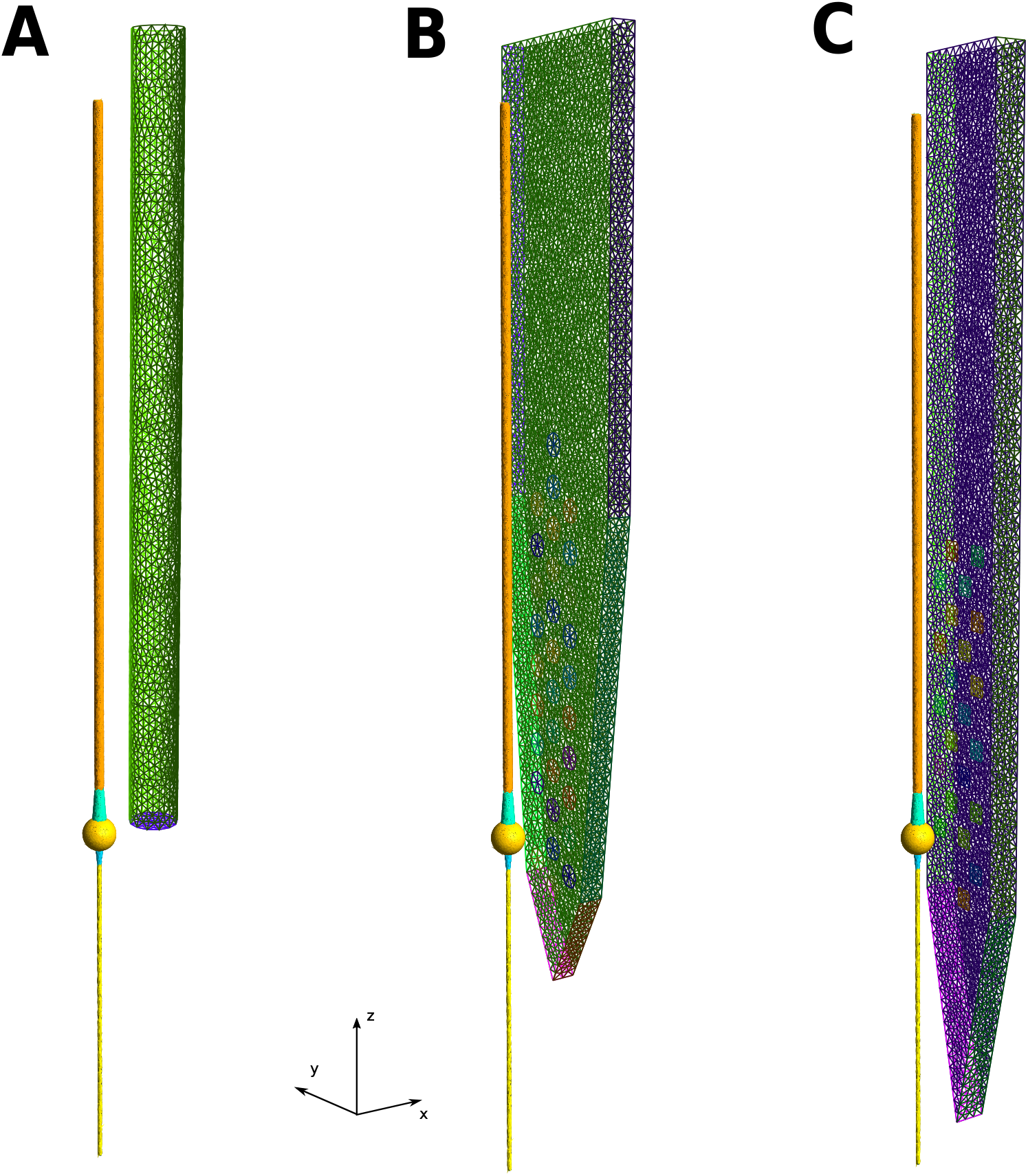
Visualization of simplified neuron and probe meshes. (A) Microwire: the probe has a 15 μm radius and it is aligned to the neuronal axis (*z* direction) and the center of the probe tip is at (40, 0, 0) μm (the soma center is at (0, 0, 0) μm). The axon and soma of the neuron are depicted in yellow, the dendrite is orange, and the axon and *dendritic* hillock are in cyan. (B) Neuronexus MEA: the probe represents a Neuronexus A1x32-Poly3-5mm-25s-177-CM32 with recording sites facing the neuron. The MEA is 15 μm thick and the center of the bottom vertex is at (40, 0, −100) μm. The maximum width of the probe is 114 μm, which makes it almost 4 times larger than the microwire probe. (C) Neuropixels MEA: this probe [35] has a width of 70 μm, a thickness of 20 μm, and the center of the bottom vertex is at (40, 0, −100) μm. All meshes represented here are built with the finest coarseness described in the text (*coarse 0*).

In order to evaluate the effect of the described probes depending on the relative distance to the neuron (*x* direction), we generated several meshes in which the distance between the contact sites and the center of the neuron was 17.5, 22.5, 27.5, 37.5, 47.5, and 77.5 μm. Note that these distances refer to the beginning of the microwire tip (which extends in the *x* direction for 30 μm and to the MEA *y* − *z* plane (for the MEA probes the recording sites do not extend in the *x* direction). When not specified, instead, the distance for the microwire probe was 25 μm, 32.5 μm for the Neuronexus MEA probe, and 30 μm for the Neuropixels probe (center of the probe tip at 40 μm).

To investigate if and how the bounding box size affects the simulation, since the electric potential is set to zero at its surface, we generated meshes with five different box sizes. Defining *dx, dy,* and *dz* as the distance between the extremity of the neuron and the box in the *x, y*, and *z* directions, the three box sizes were:

**size 1:** *d_x_* = 80 µm, *d_y_* = 80 µm, and *d_z_* = 20 µm
**size 2:** *d_x_* = 100 µm, *d_y_* = 100 µm, and *d_z_* = 40 µm
**size 3:** *d_x_* = 120 µm, *d_y_* = 120 µm, and *d_z_* = 60 µm
**size 4:** *d_x_* = 160 µm, *d_y_* = 160 µm, and *d_z_* = 100 µm
**size 5:** *d_x_* = 200 µm, *d_y_* = 200 µm, and *d_z_* = 150 µm

Moreover, we evaluated the solution convergence depending on the resolution by generating meshes with four different resolutions. Defining *r_n_*, *r_p_*, and *r_ext_* as the resolutions/typical mesh element sizes for the neuron volume and membrane, for the probe, and for the bounding box surface, respectively, the four degrees of *coarseness* were:

**coarse 0:** *r_n_* = 2 µm, *r_p_* = 5 µm, and *r_ext_* = 7:5 µm
**coarse 1:** *r_n_* = 3 µm, *r_p_* = 6 µm, and *r_ext_* = 9 µm
**coarse 2:** *r_n_* = 4 µm, *r_p_* = 8 µm, and *r_ext_* = 12 µm
**coarse 3:** *r_n_* = 4 µm, *r_p_* = 10 µm, and *r_ext_* = 15 µm

At the interface between two resolutions, the mesh size was determined as their minimum. Further, having instructed `gmsh` to not allow hanging nodes the mesh in the surroundings of the neuron and probe is gradually coarsened to *r_ext_* resolution.

For each of the mesh configuration with varying probe model, box size, and coarseness we simulated the extracellular signals with and without the probe in the extracellular space and sampled the electric potential at the recording site locations (even when the probe is absent).

### 2.3 Membrane model and finite element implementation

On the membrane of the soma and the axon, the ionic current density, *I*_ion_, is computed by the Hodgkin-Huxley model with standard parameters as given in [36]. On the membrane of the dendrite, we apply a passive membrane model with a synaptic input current of the form

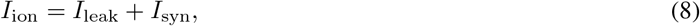

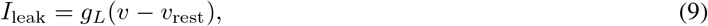

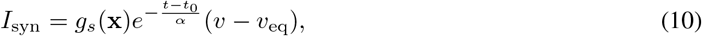

where

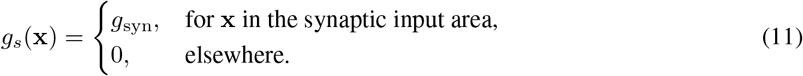

The parameters of the dendrite model are given in Table 1, and the synaptic input area is defined as a section of the dendrite of length 20 *μm* located 350 *μm* from the soma, as illustrated in Figure 1.

The EMI model (1)-(7) is solved by the operator splitting scheme and the *H*(div) discretization proposed in [19]. In this scheme a single step of the EMI model consists of two sub-steps. First, assuming the current membrane potential *v* is known, the ordinary differential equations (ODE) of the membrane model are solved yielding a new membrane state and the value of *v*. Next, (7) discretized in time with *I*_ion_ set to zero, is solved together with equations (1)-(6) using the computed value of v as input. This step yields the new values of intra/extra-cellular potentials *u_i_*, *u_e_* and the transmembrane potential *v*. The H(div) approach then means that the EMI model is transformed by introducing unknown electrical fields *σ_i_*ᐁ*u_e_* and *σ_e_*ᐁ*u_e_* in addition to the potentials *u_i_*, u_e_ and *v*. Thus more unknowns are involved, however, the formulation leads to more accurate solutions, cf. [19, section 3.].

**Table 1:**
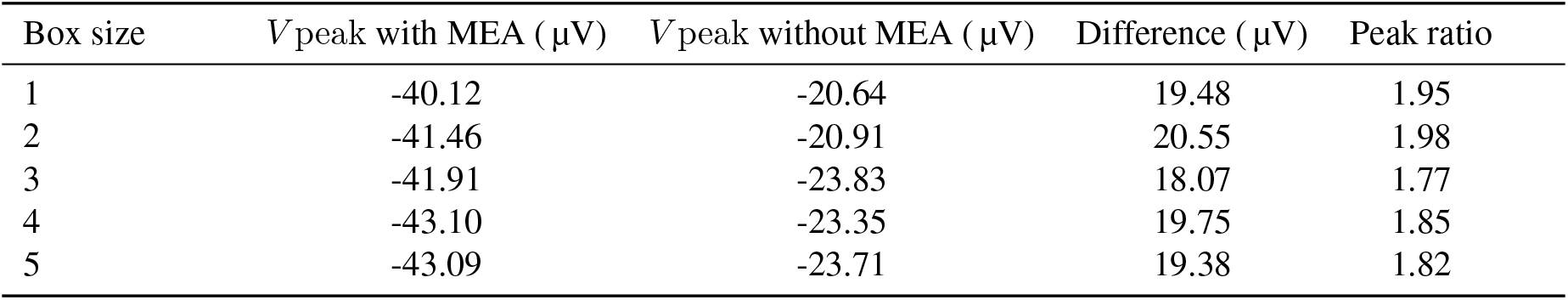
Model parameters used in the simulations. The parameters of the Hodgkin-Huxley model are given in [36].

In our implementation the ODE solver for the first step of the operator splitting scheme is implemented on top of the computational cardiac electrophysiology framework `cbc.beat` [37]. For the second step, the H(div) formulation of the EMI model, see [19, section 2.3.3], is discretized by the finite element method (FEM) using the `FEniCS` library [38]. More specifically, the electrical fields are discretized by the lowest order Raviart-Thomas elements [39] while the potentials use piecewise constant elements. The linear system due to implicit/backward-Euler temporal discretization in (7) and FEM is finally solved with the direct solver MUMPS [40] which is interfaced with `FEniCS` via the `PETSc` [41] linear algebra library.

## 2.4 Other modeling approaches

### 2.4.1 Cable equation and current summation (CS)

The cable equation [42, 43, 44] is of great importance in computational neuroscience, and it reads,

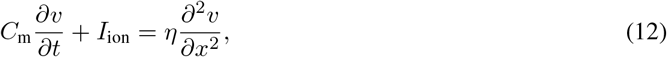

here, *v* is the membrane potential of the neuron, *C*_m_ is the membrane capacitance, *I*_ion_ is the ion current density and *η* = ^*hσ_i_*^/4, where *h* is the diameter of the neuron and *σ*_i_ denotes the intracellular conductivity of the neuron [42].

This equation is used to compute the membrane potential of a neuron and the solution is commonly obtained by dividing the neuron into compartments and replacing the continuous model (12) by a discrete model [42]. In order to compute the associated extracellular potential, it is common to use the solution of the cable equation to compute the transmembrane currents densities in every compartment, and then invoke the classical summation formula,

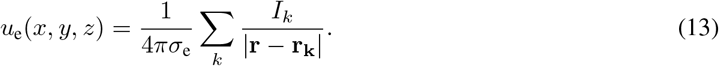

Here, *σ*_e_ is the extracellular conductivity, r_k_ is the center of the *k*-th compartment of the neuron, |r − r_k_| denotes the euclidean distance from r = r(*x, y, z*) to the point r_k_, and *I_k_* denotes the transmembrane current of each compartment. We denote this method as current summation approach (CS) [6].

We implemented the same simulations presented in Section 2.1 using the conventional modeling approach described above (CS) to compare them with the EMI simulations. We used `LFPy` [11], running upon Neuron 7.5 [9, 10], to solve the cable equation and compute extracellular potentials using Equation 13. As morphology, we used a ball-and-stick model with an axon with the same geometrical properties described in Section 2.2. Similarly to the EMI simulations, we used a synaptic input in the middle of the dendritic region activated in the EMI simulation (z=360 μm) to induce a single spike and we observed the extracellular potentials on the recording sites. The synaptic weight was adjusted so that the extracellular largest peak was coincident in time with the one from the EMI simulation. To model the spatial extent of the electrodes, we randomly drew 50 points within a recording site and we averaged the extracellular potential computed at these points [11]. We used the same parameters shown in Table 1 (note that in Neuron conductances are defined in S/cm^2^ so we set *g_L_* = *g*_pas_ = 0.06 · 10^−3^ S/cm^2^) and we used an axial resistance R_*a*_ of 150 Ω/*cm*. The `fixed_length` method was used as discretization method with a fixed length of 1 μm, yielding 658 segments (23 somatic, 422 dendritic, and 213 axonal). Transmembrane currents were considered as current point sources in their contributions to the extracellular potential, following Equation 13 (using LFPy `pointsource` argument of the `RecExtElectrode` class).

### 2.4.2 Hybrid solution (HS)

The hybrid solution (HS) [20, 21, 22] combines the transmembrane currents for each neural segments computed with the cable equation and a finite element modeling for the extracellular space. The transmembrane currents are used as source terms in a finite element solution of the Poisson Equation in the extracellular space (Equation 2, using an iterative solver for the Poisson problem, specifically, preconditioned conjugate gradients with algebraic multigrid preconditioning). With this approach, the probe can be explicitly modeled using insulating (Neumann) boundary conditions at the surface of the probe (Equation 5) and the differences between the HS and the EMI solution lie in differences regarding the modeling of the neuron dynamics, such as the self-ephaptic effect. The HS requires that a FEM simulation is run for each timestep of the transmembrane currents, each time setting the source terms with the currents at the specific timestep. This makes it computationally expensive, especially, for long simulations. Alternatively, one could run a single FEM simulation for each neural segment with a unitary test current and then use the potentials computed at the recording sites as a static map for summing the contribution of all currents at each timestep. The latter approach can be also computationally complex, as the number of segments in the multi-compartment simulation can be quite high and it would require to store in memory a large number of finite element solutions.

### 2.4.3 Method of images (MoI)

As the silicon probes are made of insulated material, they could be approximated with the method of images (MoI) [23, 24]. With the MoI the probe is assumed to be an infinite insulating plane, effectively increasing the extracellular potential by a factor of 2. Using the MoI, the factor 2 can be explained as follows: for each current source an *image* current source is introduced in the mirror position with respect to the insulating plane, effectively doubling the potential in proximity of the plane and canceling current densities normal to the plane. While the MoI uses a factor 2, assuming an insulated infinite plane, the finite-size of the probes used in this work and the results that will be shown in Section 3.1 suggest the use of a factor between slightly lower than 2 (e.g. 1.85 is used in Section 3.3.1).

### 2.4.4 Probe correction (PC)

The hybrid solution is a good and widely used approach to model a non-homogeneous extracellular space, especially in the pheripheral nervous system literature [22, 20, 21]. However, it requires to run a finite element simulation for every neuron simulation, as transmembrane currents are located in different positions for different neurons.

In order to overcome this issue, we designed the probe correction method (PC) that relies on the reciprocity principle [45]. The reciprocity principle states that if a current *I*_1_ in a position (*x*_1_, *y*_1_, *z*_1_) generates a potential *u*_1_ in a second position (*x*_2_, *y*_2_, *z*_2_), then the same current *I*_1_ placed in (*x*_2_, *y*_2_, *z*_2_) will result in a potential *u*_1_ in (*x*_1_, *y*_1_, *z*_1_)^1^. Using this principle, we first simulated with a finite element method the extracellular potential generated by a test current (1 *nA*) from each electrode *i* of a specific probe (e.g. Neuronexus) in any point of the extracellular space and define it as *u_i_*(*x_i_*, *y_i_*, *z_i_*), where (*x_i_*, *y_i_*, *z_i_*) is the relative position with respect to the electrode *i*. Also in this case we used an iterative solver for the Poisson problem (preconditioned conjugate gradients with algebraic multigrid preconditioning). Then, leveraging on the reciprocity principle, we mapped the contribution of each transmembrane current to the potential at each electrode *i* as: *u_ik_* = *I_k_u_i_*(*x_k_*, *y_k_*, *z_k_*), where (*x_k_*, *y_k_*, *z_k_*) is now the relative position between the *k*-th neural segment and the electrode *i*, and *I_k_* is the transmembrane current for the *k*-th neural segment. The potential at each electrode *i* can be computed as:

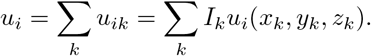

The PC method allows to pre-compute the effect of a probe in the extracellular space and then use this mapping for any neural model, without the need to run a full FEM simulation. The number of FEM solution that need to be computed and stored during the pre-mapping is equal to the number of electrodes in the probe.

## 3 Results

In this section we present results of numerical simulation which quantify the effect of introducing probes in the extracellular domain on the extracellular potential. We show how this effect depends on the distance between the neuron and the probe, their lateral alignment, and the probe rotation. Furthermore, we evaluate the numerical variability of the solutions, we compare with other modeling schemes, and finally report CPU-efforts for the simulations.

### 3.1 The *probe effect*

#### 3.1.1 The geometry of the probe affects the recorded signals

The first question that we investigated is whether the probes have an effect and, if so, how substantial this effect is and if it depends on the probe geometry. In order to do so we analyzed the extracellular action potential (EAP) traces with and without placing the probe in the mesh.

In Figure 3 we show the EAP with and without the microwire probe (A), the Neuronexus probe (B), and the Neuropixels probe (C). The blue traces are the extracellular potentials computed at the recording sites when the probe was removed, while the orange traces show the potential when the probe is present in the extracellular space. In this case the probe tip was placed 40 μm from the soma center, we used a box of size 2 and coarse 2 resolution. It is clear that the *probe effect* is more prevalent for the MEA probes than for the microwire, suggesting that the physical size and geometry of the probe plays an important role. In particular, for the Neuronexus probe the minimum peak without the probe is −21.09 μV and with the probe it is −41.26 μV: the difference is 20.17μV. For the Neuropixels probe the peak with no probe is −21.2 μV, with the probe it is −44.36 μV and the difference is 23.16 μV. In case of the microwire type of probe, the effect is minimal: the minimum peak without the probe is −16.85 μV, with the probe it is −15.82 μV, and the difference is about 1. 03 μV (the peak without the probe is even larger than the one with the probe). Note that the values for the microwire are slightly lower than the MEAs because even if the microwire tip center is at the same distance (40 μm), it extends for 30 μm in the *x* direction, effectively lowering the recorded potential due to the fast decay of the extracellular potential with distance. The recording sites of the MEAs, instead, lie on the *y* − *z* plane, at a fixed distance.

**Figure 3:**
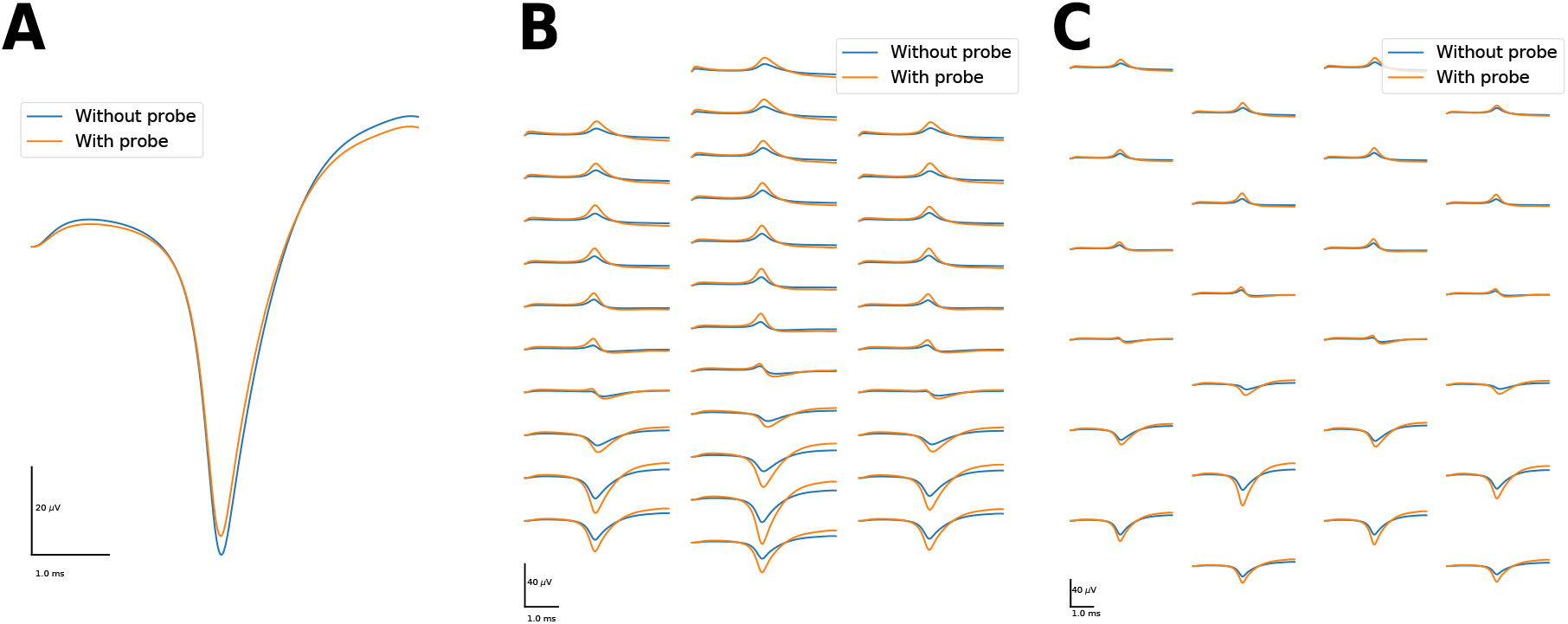
Extracellular action potentials (EAPs). (A) EAPs without (blue) and with (orange) the microwire probe (single recording site) in the extracellular space. The amplitude difference in the largest peak is only 1.3 μV, which is negligible for most applications. (B) Same as (A) but with the Neuronexus MEA probe. For this probe, the difference in amplitude is 20.17 μV (the solution with the MEA is almost twice as large as the one without the MEA in the extracellular space). (C) Same as (A) but with the Neuropixels MEA probe. For this probe, the difference in amplitude is 23.16 μV.

The MEAs, electrically speaking, are like insulating *walls* that do not allow currents to flow in. The insulating effect can be appreciated in Figure 4, in which the extracellular potential at the time of the peak is computed in the [10, 100] μm interval in the *x* direction and in the [−200, 200] μm interval in the *z* direction (the origin is the center of the soma). Panel A shows the extracellular potential with the probe (Neuronexus) and panel B without the probe. The currents are deflected due to the presence of the probe, and this causes an increase (in absolute value) in the extracellular potential between the neuron and the probe, as shown in panel C, where the difference of the extracellular potential with and without probe is depicted. The substantial effect using the MEA probe probably also depends on the arrangement of the recording sites: while for the MEAs, the electrodes *face* the neuron (they lie on the *y* − *z* plane) and currents emitted by the membrane cannot flow in the *x* direction due to the presence of the probe, for the microwire, the electrode is at the tip of the probe (at *z* = 0, extending in the *x* − *plane* – see Figure 2) and currents can *naturally* flow downwards in the *x* direction, yielding a little effect (Figure 4C shows that the effect at the tip of the MEA probe is almost null).

**Figure 4:**
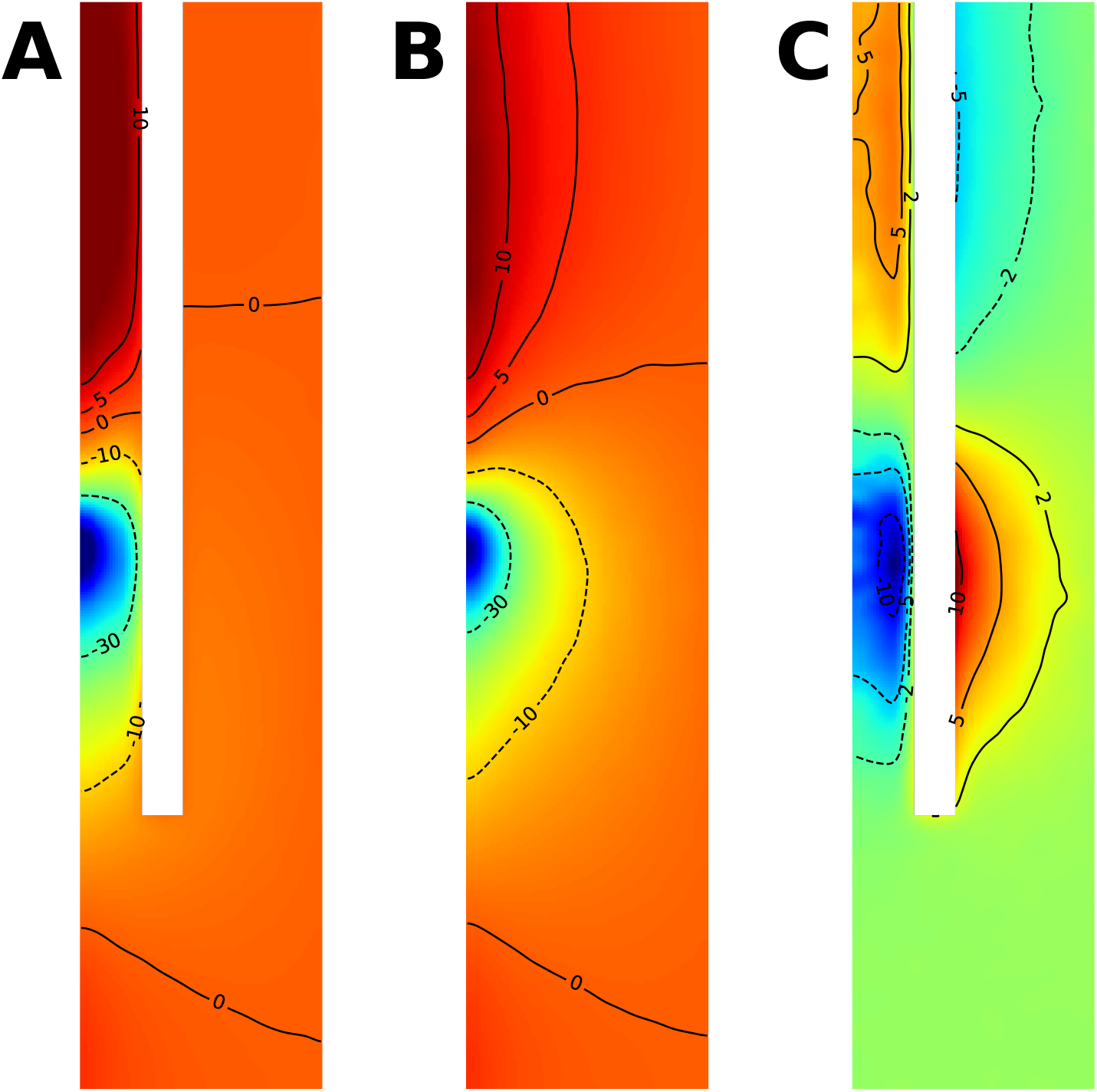
Extracellular potential distribution on the *x* − *z* plane with the Neuronexus MEA probe (A) without the probe (B), and their difference (C). The images were smoothed with a gaussian filter with standard deviation of 4 μm. The color code for panel A and B is the same. The isopotential lines show the potential in μV. The probe (white area) acts as an insulator, effectively increasing the extracellular potential (in absolute value) in the area between the neuron and the probe (panel C, blue colors close to the soma and red close to the dendrite) and decreasing it behind the probe of several μV. The effect is smaller at the tip of the probe (the green color represents a 0 μV difference).

#### 3.1.2 The amplitude ratio is constant with probe distance

In this section we analyze the trend of the probe-induced error depending on the vicinity of the probe. We swept the extracellular space from a closest distance between the probe and the somatic membrane of 7.5 μm to a maximum distance of 67.5 μm.

In Figure 5A-B-C we plot the absolute peak values with (orange) and without probe (blue), as well as their difference (green) for the microwire (A), Neuronexus (B) and Neuropixels (C) probes. For the microwire (A), as observed in the previous section, the *probe effect* is small and the maximum difference is 1.97 μV, which is 10.1 % of the amplitude without probe, when the probe is closest. For the Neuronexus MEA probe (B), at short distances the difference between the peaks with and without probe is large − 40.5 μV (88.8 % of the amplitude without probe) at 7.5 μm probe-membrane distance – and it decreases as the probe distance increases. At the farthest distance, where the probe is at 72.5 μm from the somatic membrane, the difference is 4.38 μV, which is 90.2 % of the amplitude without probe. For the Neuropixels MEA probe (C) the effect is in line with the Neuronexus probe, with a maximum difference of 41.07 μV (95.9 % of the amplitude without probe) when the probe is closest and a minimum of 5.08 μV, which is still 116.1 % of the amplitude without probe, when the probe is located at the maximum distance. Note that the peak amplitudes on the microwire probe are smaller than the one measured on the MEAs at a similar distances. At the closest distance, for example, the Neuronexus MEA electrodes lie on the *y* − *z* plane exactly at 7.5 μm from the somatic membrane. For the microwire, instead, 7.5 μm is the distance to the beginning of the cylindrical probe, whose tip extends in the *x* direction for 30 μm. The simulated electric potential is the average of the electric potential computed on the microwire tip and it results in a much lower amplitude due to the fast decay of the extracellular potential with distance (see Equation 13).

**Figure 5:**
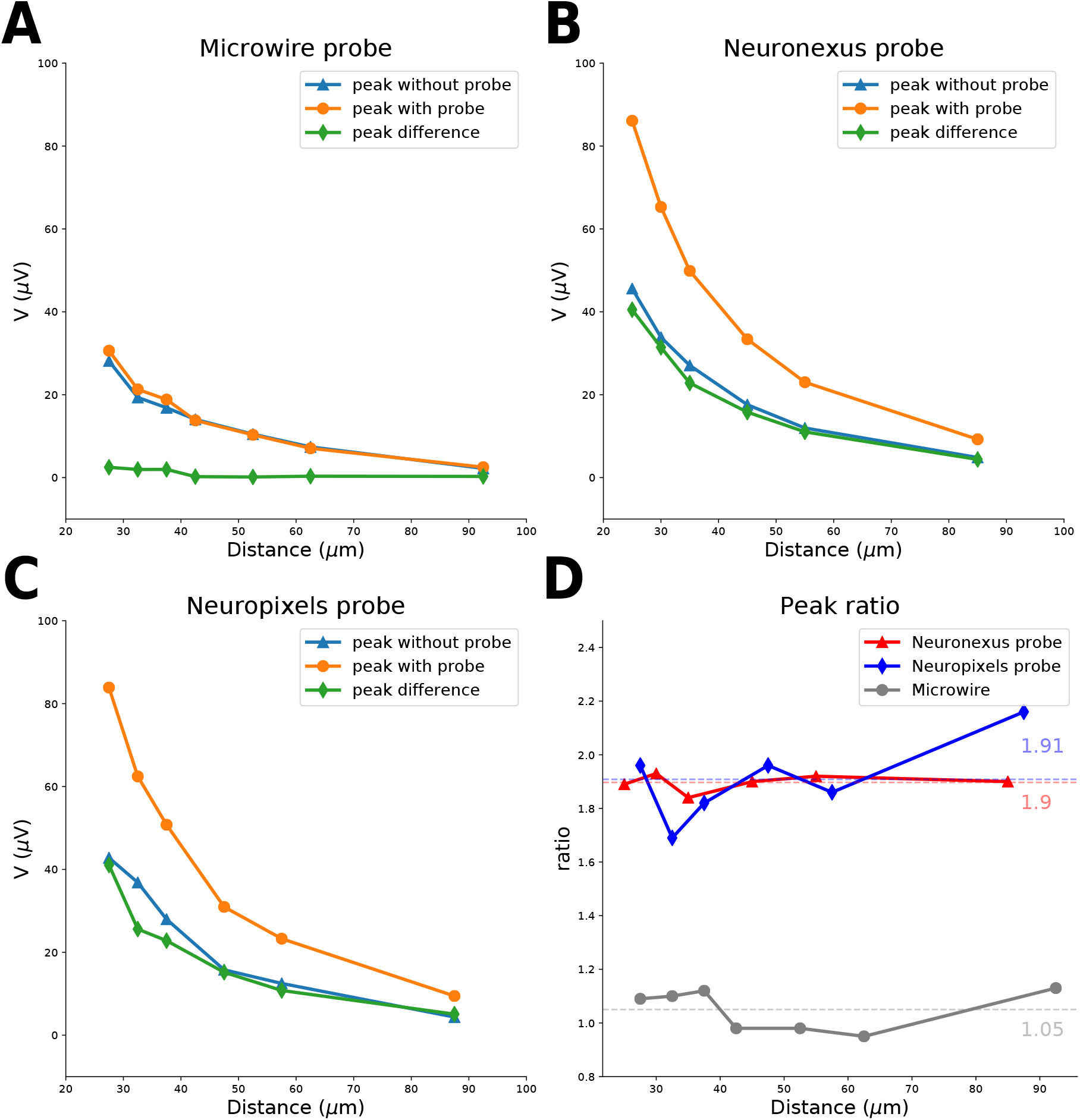
Differences in EAP maximum absolute value peak with and without probe depending on distance. (A) Microwire probe: maximum peak without probe (blue), with probe (orange), and their difference (green). The difference is small even when the probe is close to the neuron. (B) Neuronexus MEA probe: maximum peak without probe (blue), with probe (orange), and their difference (green). The difference is large at short distances and it decays at larger distances. (C) Neuropixels MEA probe: maximum peak without probe (blue), with probe (orange), and their difference (green). Also for this probe the difference is large at short distances and it reduces at further away from the neuron. (D) Ratio between peak with and without probe for Neuronexus (red), the Neuropixels (blue) and the microwire probe (grey). The ratio is almost constant at different distances and the average value is 1.9 for the Neuronexus, 1.91 for the Neuropixels, and 1.05 for the microwire probe.

In panel D of Figure 5 we show the ratio between the peak with probe and without probe depending on the probe distance for the Neuronexus (red), Neuropixels (blue), and the microwire (grey) probes. The ratio for the microwire probe varies around 1 (average=1.05), confirming that the *probe effect* can be neglected for microwire-like types of probe, due to their size and geometry. Instead, when a MEA probe is used, the average ratio is around 1.9 and its effect on the recordings cannot be neglected.

#### 3.1.3 The probe effect is reduced when neuron and probe are not aligned

So far, we have shown results in which the neuron and the probe are perfectly aligned in the y direction, but the *probe effect* is likely to be affected by the neuron-probe alignment, since the area of the MEA probe (we focus here on the Neuronexus and Neuropixels MEA probes as the effect using the microwire is negligible) *facing* the neuron changes depending on the lateral shift in the *y* direction and probe rotation.

To quantify the trend of the *probe effect* depending on the y shift, we ran simulations moving the probes at different y locations (10, 20, 30, 40, 50, 60, 80, and 100 μm) and computed the ratios between the maximum peak with and without the MEA in the extracellular space. The simulations were run with *coarse 2* resolution and *boxsize 5* and the probe tip was at 40 μm from the center of the neuron. In Figure 6A we show the peak ratios depending on lateral y shifts. The ratio appears to decrease almost linearly with the shifts, from a value of around 1.8–1.9 when the probe is centered (note that the peak ratio slightly varies depending on resolution and size, as covered in Section 3.2) to a value of around 1.2 when the shift is 100 μm (the half width of the probe is 57 μm for Neuronexus and 35 μm for Neuropixels).

**Figure 6:**
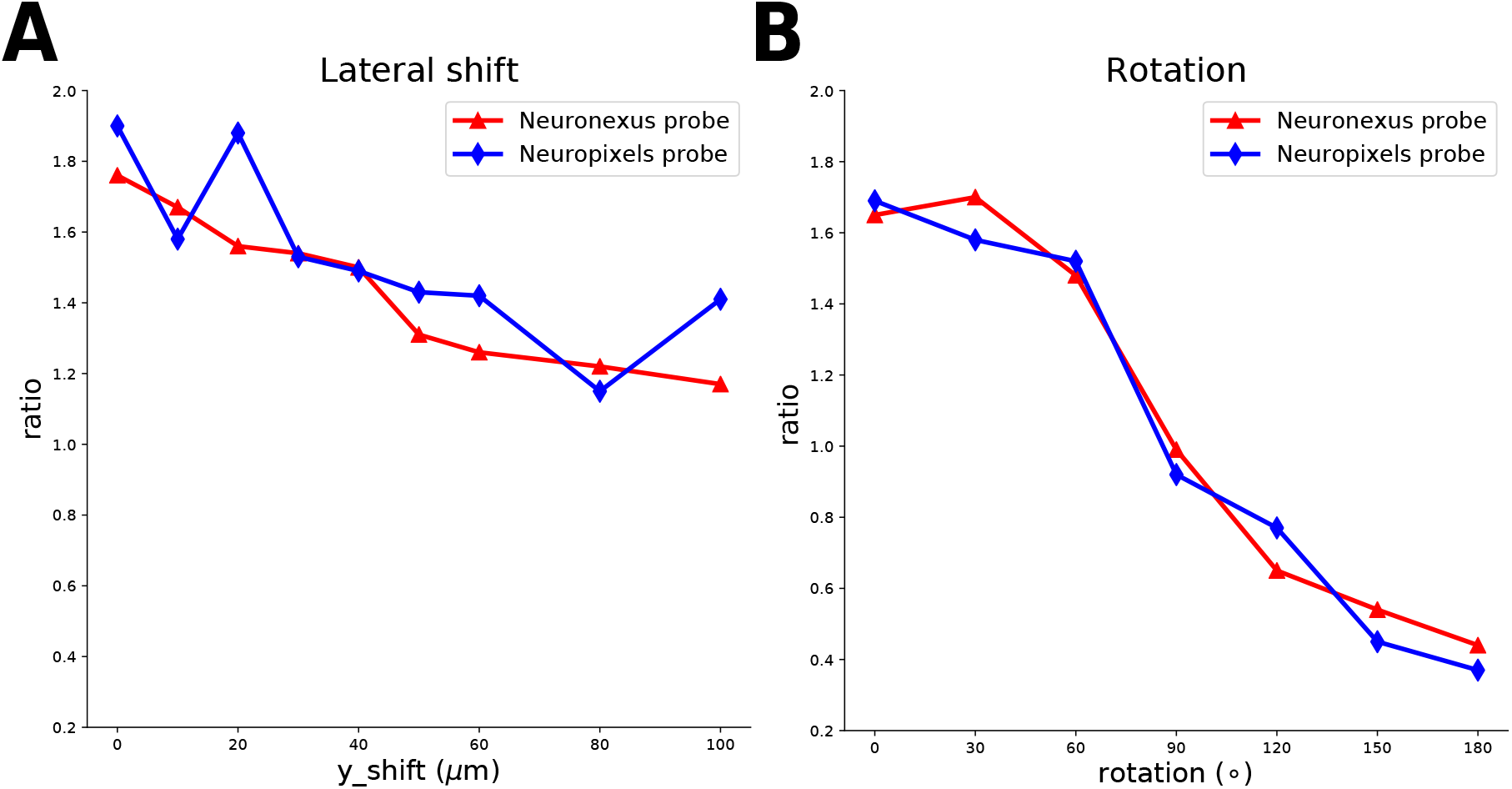
Effects of neuron probe alignment. (A) Amplitude ratio for different y lateral shifts for the Neuronexus (red) and Neuropixels (blue) probes. The ratio is decreases almost linearly with the y shift. (B) Amplitude ratio for different probe rotations for the Neuronexus (red) and Neuropixels (blue) probes. At small rotations, the peak ratio is between 1.6 and 1.8, at 90° rotation (when the probe exposes its thinnest side to the neuron) it is around 1, and between 90° and 180° the shadowing effect of the probe makes the ratio lower than 1.

In order to evaluate the effect of rotating the probes, we ran simulations with the probe at 70 μm distance (to accommodate for different rotations), *coarse 2* resolution, *boxsize 4,* and rotations of 0, 30, 60, 90, 120, 150, and 180°. In Figure 6B the peak ratios depending on the rotation angle are shown. For small or no rotations (0, 30°) the value is around 1.7 (note that we always selected the electrode with the largest amplitude, which might not be the same electrode for all rotations). For a rotation of 90° the peak ratio is around 1 (the probe exposes its thinnest side to the neuron) and for further rotations the probe’s shadowing effect makes the peak with the probe smaller (as observed in Figure 4C), yielding peak ratio values below 1. These results demonstrate that the relative arrangement between the neuron and the probe play an important role in affecting the recorded signals.

### 3.2 EMI solution dependence on domain size and resolution

We generated meshes of four different resolutions and five different box sizes, as described in Section 2.2, in order to investigate how the resolution and the domain size affect the finite element solutions. Since we are mainly interested in how the probe affects the extracellular potential and we showed that only for MEA probes this effect is large, we focus on the extracellular potential at the recording site with the maximum negative peak. We used the Neuronexus MEA probe for this analysis and the distance of the tip of the probe was 40 μm (the recording sites plane is at 32.5 μm from the somatic center). The recording site which experienced the largest potential deflection was at position (32.5,0, −13) μm, i.e. the closest to the neuron soma in the axon direction. For a deeper examination of convergence of the EMI model refer to [6]. For resolutions *coarse 0* and *coarse 1* the box of size 4 and 5, and of size 5, respectively, were too large to be simulated.

In Table 2 we show the values of the minimum EAP peak with and without the Neuronexus probe, their difference, and their ratio grouped by the domain (box) size and averaged over resolution. Despite some variability due to the numerical solution of the problem, there is a common trend in the peak values as the domain size increases: the minimum peaks tend to be larger in absolute values, both when the probe is in the extracellular space (from −40.12 μV for box size 1 to −43.09 μV for box size 5) and when it is not (from −20.64 μV for box size 1 to −23.71 μV for box size 5). This can be explained by the boundary conditions that we defined for the bounding box (Equation 3), which forces the electric potential at the boundaries to be 0. For this reason, a smaller domain size causes a steeper reduction of the extracellular potential from the neuron to the bounding box, making the peak amplitude, in absolute terms, smaller. The peak difference with and without the MEA probe appears to be relatively constant, but the peak ratio tends to slightly decrease with increasing domain size for the same reason expressed before (from 1.95 for box size 1 to 1.82 for box size 5). The solutions appear to be converging for box sizes 4 and 5, but the relative error (difference between box 1 and box 5 values divided by the value of box 5) is moderate (6.89 % for the peak with probe, 12.95 % for the peak without probe, and 4.14 % for the peak ratio). Nevertheless, the 1.8–1.85 peak ratio values obtained with larger domain sizes should be a closer estimate of the true value.

**Table 2:**
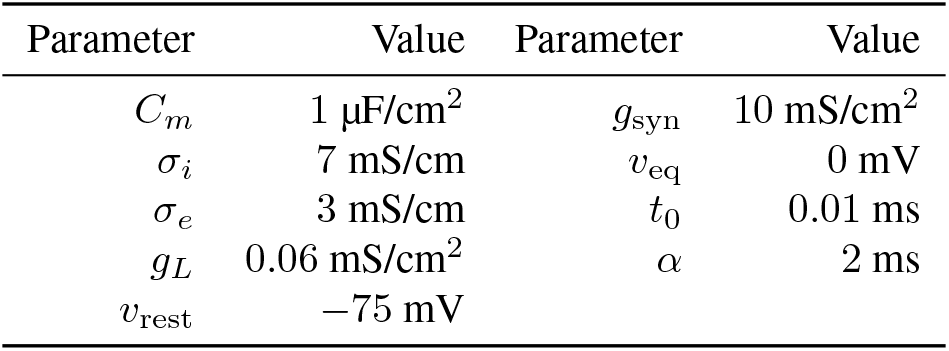
Solution variability depending on box (domain) size. The columns contain the maximum peak with the Neuronexus (MEA) probe, without the probe, the difference and ratio of the amplitudes with and without probe. The values are averaged over all resolutions.

Table 3 displays the same values of Table 2, but with a fixed box size of 2 and varying resolution (*Coarseness*). The relative error (maximum difference across resolutions divided by the average values among resolutions) of the peak with the MEA is 3.3 %, without the probe it is 6.65 %, and for the peak ratio it is 3.53 %.

**Table 3:** Solution variability depending on resolution (*Coarseness*). The columns contain the maximum peak with the Neuronexus (MEA) probe, without the probe, the difference and ratio of the amplitudes with and without probe. The values are computed with a box size 2.

| Coarseness | $V_{\text{peak with MEA}} (\mu\text{V})$ | $V_{\text{peak without MEA}} (\mu\text{V})$ | Difference ($\mu\text{V}$) | Peak ratio |
| --- | --- | --- | --- | --- |
| 0 | -41.74 | -20.67 | 21.07 | 2.02 |
| 1 | -40.74 | -20.25 | 20.49 | 2.01 |
| 2 | -41.26 | -21.09 | 20.18 | 1.96 |
| 3 | -42.11 | -21.64 | 20.46 | 1.95 |

Because the main purpose of this work was to qualitatively investigate the effect of various probe designs and the effect of distance, alignment, and rotation on the measurements, we used resolution *coarse 2* and box size 2, which represented an acceptable compromise between accuracy and simulation time. For investigating the effect of probe rotation and side shift we increased the box size to 4 and 5, respectively, to accommodate the position of the neural probe. Finally, in Section 3.3 we increased the resolution to *coarse 0* and used box size 3 to obtain more accurate results for the comparison with the cable equation simulations.

### 3.3 Comparison with other approaches

After having investigated how an extracellular probe affects the amplitude of the recorded potentials and how this amplitude is modulated with distance, alignment, and rotation between the neuron and the probe, we now compare the EMI solution to other modeling approaches. We first analyze the differences between the EMI solution without the probe and the cable equation / current summation approach (CS) and between the EMI solution with the probe and the hybrid solution (HS). Then we focus on the HS, which combines a cable equation solution and an explicit model of the extracellular space, including the probe, in a FEM framework, and compare its solution to two correction strategies: the method of images (MoI) and the probe correction (PC).

In all the following simulations we used a mesh with *coarse 0* resolution and box size 3. The distance between the neuron soma center and the probe tip was 40 μm, resulting in recording sites on the *x* = 32.5 μm plane.

#### 3.3.1 EMI, CS, and HS comparison

In order to compare the EMI simulations to conventional modeling, we built the same scenario shown in Figure 2B (Neuronexus probe) using Neuron and LFPy, as described in Section 2.4. As conventional modeling assumes an infinite and homogeneous medium, we compared the EAPs obtained combining the cable equation solution (Equation 12) and the current summation formula (Equation 13) with the EMI simulations without the probe. The extracellular traces for the current summation approach (CS, red) and the EMI model (blue) are shown in Figure 7A. The EAPs almost overlap for every recording site, despite some differences in amplitude. On the electrode with the largest peak, the value for the EMI solution is −23.03 μV, while the value for the CS is −27.95 μV (the difference is 4.91 μV). This difference, which has been previously observed, is intrinsic to the EMI model [6], and can be due to self-ephaptic effects [12, 13, 14, 15, 16, 17, 6]. Note also that the condition that forces the extracellular potential to zero at the boundary of the domain causes a steeper descent in the extracellular amplitudes, as discussed in Section 3.2.

**Figure 7:**
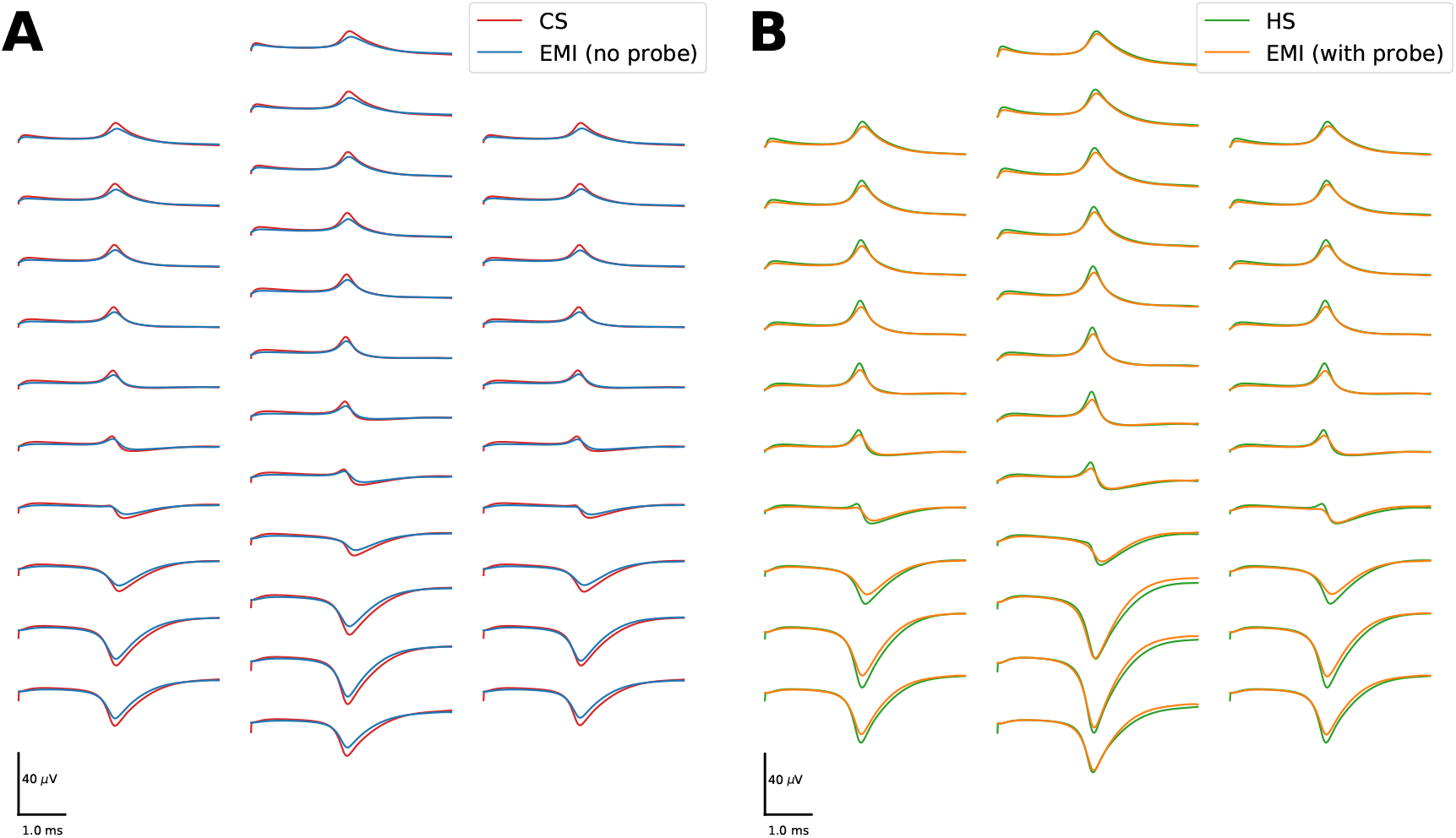
Comparison of the EAPs (A) between the current summation approach (CS, red) and the EMI model without probe (blue), displaying a peak amplitude difference of 4.91 μV, and (B) between the hybrid solution (HS, green) and the EMI model with probe (orange), exhibiting a peak amplitude difference of 3.55 μV.

The hybrid solution (HS) uses currents computed with the cable equation and runs a FEM simulation of the extracellular space, including the probe. In Figure 7B we show the extracellular potential of the EMI simulation with probe (orange) and the HS (green). Also in this case we observe that the EMI solution yields slightly smaller amplitudes with respect to the HS (EMI peak: −42.6 μV; HS peak −46.15 μV; difference: 3.55 μV) and these differences can be once again traced back to underlying differences of the neural solver.

#### 3.3.2 HS, MoI, and PC comparison

After having shown that there are intrinsic differences between the EMI model and solutions based on the cable equation (CS, HS), we now compare two computationally less expensive strategies that could be used to account for the *probe effect* in modeling of extracellular potentials.

The MoI is an attractive candidate due to its almost null computational cost, as it only multiplies all values by a constant factor. The factor for infinite insulated planes, as described in Section 2.4.3, is 2, but as shown in Figures 5 and 6, for MEA probes it is somewhere between 0 and 2 depending on the neuron-probe lateral shift and rotation. In this scenario, the neuron is perfectly aligned with the probe and there is no rotation. The peak ratio was computed by dividing the peaks of the EMI solution with and without the probe and it was set to 1.85. In Figure 8A the EAP from the HS (green), from the MoI with factor 2 (pink), and from the MoI with factor 1.85 (grey) are displayed. The MoI (pink) overshoots the estimation of the extracellular amplitudes (MoI peak: −55.89 μV; HS −46.15 μV; difference: 9.74 μV). Even when adjusting the MoI amplitude from the findings obtained from the EMI simulations, the amplitudes are slightly larger than the HS (1.85 MoI peak: −51.7μV; HS −46.15 μV; difference: 5.55 μV). Figure 8B shows the distribution of peak ratios of all the 32 electrodes with respect to the HS peaks. The CS and MoI solutions (including the 1.85 MoI) display a range of values in the peak ratios, showing that the amplitude modulation of the electrodes is not a constant value. This can be traced back to the fact that a lateral shift of the neuron reduces the peak ratio (Figure 6A): electrodes on the side of the probe yield a lower effect than the ones at the center of the probe. Due to this variability, a correction strategy based on the MoI will not be able to accommodate for this effect, as it multiplies the potential by a constant value.

**Figure 8:**
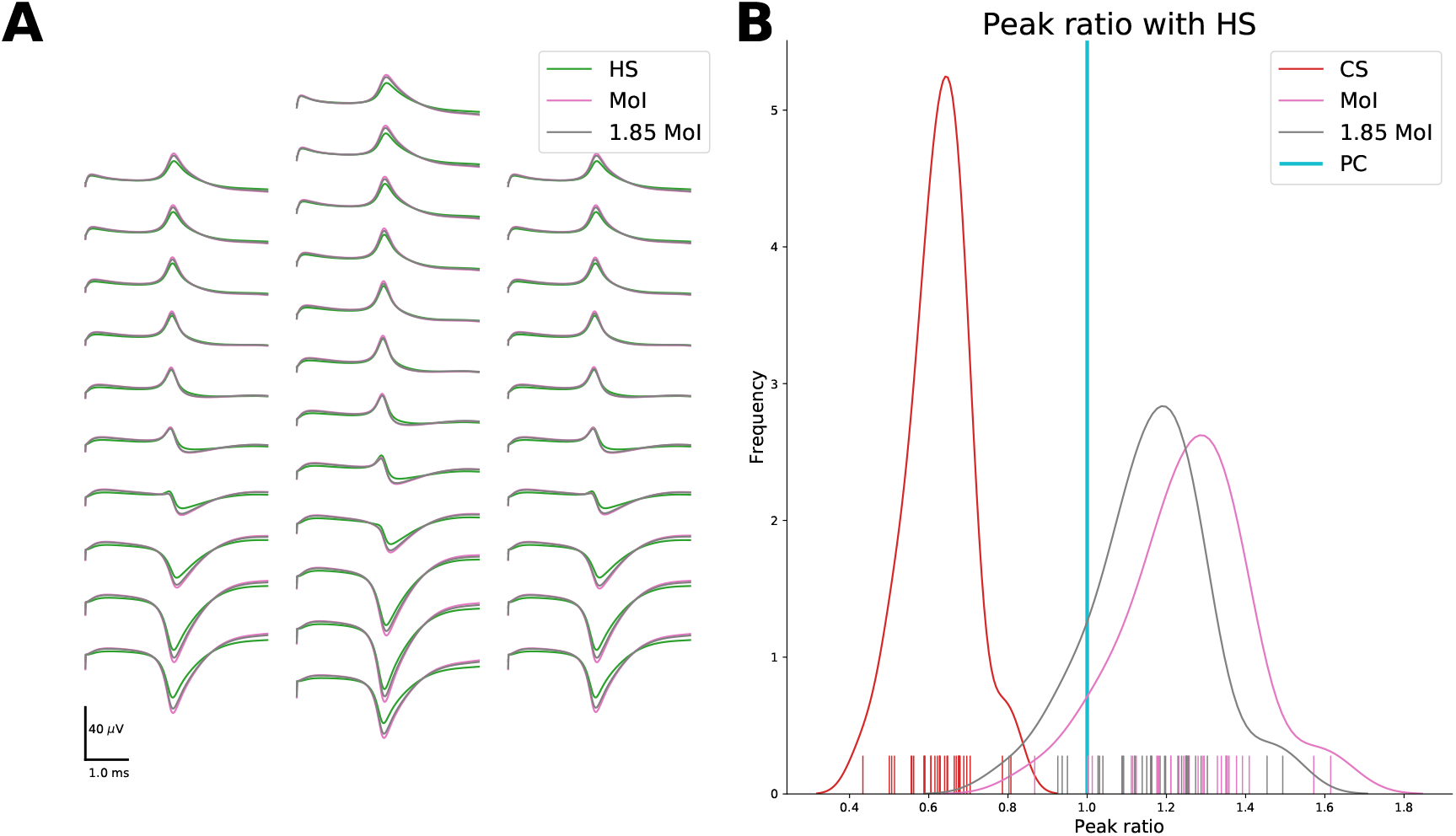
(A) EAPs of the Neuronexus probe as computed using the hybrid solution (HS, green), the method of Images (MoI, pink) and the method of Images with factor 1.85 (1.85 MoI, grey). (B) Peak ratio distribution of the electrodes of the Neuronexus probe compared to the hybrid solution, from the current summation (CS, red), method of Images (MoI, pink), the method of Images with factor 1.85 (1.85 MoI, grey), and the probe correction (PC, cyan) models. Note that the peak amplitudes computed from all the electrodes by the PC and HS approaches overlap perfectly, thus resulting in a single vertical line at peak ratio value 1.

The probe correction (PC) solution, based on the reciprocity principle (Section 2.4.4), results in a solution perfectly coincident to the HS, at a much smaller computational cost (see Table 4). In Figure 8B the PC ratios are depicted as a vertical line at 1 because the peak amplitudes are exactly the same as the HS. The PC approach, in fact, pre-maps the effect of each electrode on the extracellular domain, effectively modeling in an efficient way the distribution of peak ratios observed when using the MoI-based methods.

**Table 4:** Model type, FEM system size, resolution (*Coarseness*), box size, mesh parameters (number of cells, number of facets, number of neuron facets, and vertices), and CPU time to solve the EMI model with no probe in the extracellular space for different resolutions (*Coarseness*) and domain sizes (*Box size*). Note that for *coarse 2* and *coarse 3* the resolution of the neuron (*r_n_* = 4 μm) is the same.

| Model | System size | Coarse | Box size | Mesh Cells | Total facets | Neuron facets | Vertices | T (s) |
| --- | --- | --- | --- | --- | --- | --- | --- | --- |
| EMI | 337 515 | 3 | 1 | 66171 | 135672 | 2552 | 12400 | 1414.24 |
| EMI | 516 079 | 3 | 2 | 101443 | 207318 | 2552 | 18628 | 2813.22 |
| EMI | 562 137 | 2 | 1 | 110363 | 225887 | 2480 | 20420 | 2589.83 |
| EMI | 745 789 | 3 | 3 | 146905 | 299442 | 2552 | 26540 | 4569.11 |
| EMI | 835 365 | 2 | 2 | 164331 | 335517 | 2480 | 29940 | 4753.39 |
| EMI | 1 204 001 | 2 | 3 | 237259 | 483371 | 2480 | 42666 | 10797.78 |
| EMI | 1 225 082 | 1 | 1 | 241402 | 491840 | 3888 | 43373 | 9593.98 |
| EMI | 1 254 096 | 3 | 4 | 247514 | 503291 | 2552 | 44013 | 10756.46 |
| EMI | 1 881 777 | 1 | 2 | 371471 | 755153 | 3888 | 65867 | 18880.78 |
| EMI | 1 983 058 | 3 | 5 | 391986 | 795536 | 2552 | 68875 | 23756.09 |
| EMI | 2 110 421 | 2 | 4 | 416949 | 846736 | 2480 | 73535 | 21582.90 |
| EMI | 2 532 813 | 0 | 1 | 501235 | 1015789 | 8376 | 87535 | 27676.91 |
| EMI | 2 728 288 | 1 | 3 | 539518 | 1094385 | 3888 | 94417 | 45430.64 |
| EMI | 3 486 058 | 2 | 5 | 689996 | 1398031 | 2480 | 119968 | 53132.75 |
| EMI | 3 810 512 | 0 | 2 | 755076 | 1527718 | 8376 | 130389 | 52495.76 |
| EMI | 4 802 239 | 1 | 4 | 951245 | 1925497 | 3888 | 164359 | 68474.42 |
| EMI | 5 271 370 | 0 | 3 | 1045440 | 2112965 | 8376 | 179195 | 82601.74 |
| HS | 403 085 | 0 | 3 | 2299046 | 4665105 | - | 403085 | 3572.91 |
| PC (map) | 403 085 | 0 | 3 | 2299046 | 4665105 | - | 403085 | 2015.02 |
| PC (load) | 403 085 | 0 | 3 | 2299046 | 4665105 | - | 403085 | 409.91 |
| PC (run) | 403 085 | 0 | 3 | 2299046 | 4665105 | - | 403085 | 3.51 |

## 3.4 CPU requirements

Whereas the EMI formulation represents a powerful and more detailed computational framework for neurophysiology simulations, it is associated with a much larger computational load. The simulations were performed on an Intel(R) Xeon(R) CPU E5-2623 v4 @ 2.60GHz machine with 16 cores and 377 GB RAM running Ubuntu 16.04.3 LTS.

Table 4 contains the coarseness, domain size, number of tetrahedral cells, number of mesh vertices, total number of triangular cells (facets), facets on the surface of the neuron, the system size for the FEM problem, and the time in second (CPU time) to compute the solution for meshes without the probe in the extracellular domain. We show the results without probes in the extracellular domain, as they are they are computationally more intense due to the fact that the volume inside the probe is not meshed (although the resolution on the probe surface is finer, the resulting system size without the probe is larger than with the probe). The CPU requirements and the time needed to run the simulation strongly depends on the resolution of the mesh: the problem with coarseness 3 and box size 3 takes around 1 hour and 20 minutes (system size=745 789), while for the same box size and coarseness 0, the time required is around 22 hours (system size=5 271 370). The domain size also strongly affects the mesh size and computation time. For example, for the *coarse 2* resolution, with respect to *box 1, box2* is 1.83x slower, *box3* 4.16x, *box4* 8.33x, *box5* 20.51x.

The last four rows shows the CPU requirements for the HS and the different steps of the PC solution. These simulations, despite having the same resolution and box size as the most intense EMI simulation (*coarse 0* and box size 3), result in a much smaller system size, as they solve for the extracellular potential only (EMI also solves for intracellular potentials and currents in the entire domain). To perform a fair comparison with the EMI model, the computations were done in serial. Parallel solvers would likely speed up the HS and PC solutions and could be easily implemented. Simulating 5 ms using the HS takes about one hour, compared to the 22 hours of the EMI solution. The PC performance is divided in three steps. *PC* (*map*) refers to the the computation of the 32 FEM solutions (one for each Neuronexus solutions), and it takes slightly more than 30 minutes. Once the pre-map is computed it can be used for any neural model. Loading the FEM solutions in memory (*PC* (*load*)) requires around 7 minutes and once loaded, it takes a few seconds (3.51 s) to compute the extracellular potential. While the HS and EMI solutions computation time increase with the duration of the simulation linearly, as they iteratively solve each timestep, the PC solution multiplies each transmembrane current timeseries for a pre-defined mapping. When we ran a 500 ms Neuron simulation and then computed the extracellular potentials with the PC method the *PC* (*run*) step took only 5.38 s.

## 4 Discussion

In this article, we have used a detailed modeling framework – the Extracellular-Membrane-Intracellular (EMI) model [6,19] – to evaluate the effect of placing an extracellular recording device (neural probe) on the measured signals. We used meshes representing a simplified neuron and two different kind of probes: a microwire (a cylindrical probe with diameter of 30 μm) and Multi-Electrode Arrays (MEAs), modeling a Neuronexus commercially available silicon probe and the Neuropixels probe [35]. We quantified the *probe effect* by simulating the domain with and without the probe in the extracellular domain and we showed that the effect is substantial for the MEA probes (Figure 3B-C), while it is negligible for microwires (Figure 3A). The amplitude of the largest peak using the MEA probes is almost twice as large (~ 1.9 times) compared to the case with no probe, and this factor is relatively independent of the probe distance (Figure 5D), but it is reduced when the neuron and the probe are shifted laterally (Figure 6A) or when the probe is rotated (Figure 6B). Moreover, we discussed the effect of varying the mesh resolution and of the size of the computational domain. We also compared our finite element solutions to solutions obtained by solving the conventional cable equation, and found that the latter gave result very similar to the finite element solution when the probe was removed from the extracellular space (Figure 7A). Therefore, we suggest that the *probe effect* can be a key element in modeling experimental data obtained with MEA probes. However, clearly further analysis is needed to clarify this matter. At present the computational cost of the EMI model prevents simulations of neurons represented using realistic geometries. Thus, in an effort to offer less computationally expensive solutions to include the *probe effect* in simulations, we investigated various correction methods resulting in more accurate predictions and we proposed the probe correction method, which allows to obtain accurate solutions with reasonable computational cost and resources.

### 4.1 Comparison with previous work

In this work we used a finite element approach [19] to simulate the dynamics of a simplified neuron and to compute extracellular potentials using the EMI model. The use of FEM modeling for neural simulations has been performed before [28, 46, 47, 48, 29], but mainly as an advanced tool to study neural dynamics and ephaptic effects. In *Moffit et al.* [46], the authors simulated, using the cable equation approach, a neuron at 65 μm from a shank microelectrode with a single recording site, and then used the currents in a finite element implementation of the extracellular domain, including the shank microelectrode. They found that the amplitude of the recorded potential with the shank was 77–100 % larger than the analytical solution, but the spike shape was similar to the analytical solution (Equation 13), in accordance with our results (Figures 7A-B). The effects using MEA probes and varying distances, lateral shifts, and probe rotations were not investigated. In *Ness et al.* [24], an analytical framework for *in-vitro* planar MEA using the method of images [23] was developed. A detailed neural model was simulated using the cable equation and transmembrane currents were used as forcing functions for a finite element simulation to validate the analytical solutions. In the *in-vitro* case, in which the MEA is assumed to be an infinite insulating plane, the authors showed that the insulating MEA layer affects the amplitudes of the recorded potentials, effectively increasing it by a maximum factor of 2, which can be analytically predicted by the method of images (MoI). Using the MoI, the factor 2 can be explained as follows: for each current source an *image* current source is introduced in the mirror position with respect to the insulating plane, effectively doubling the potential in proximity of the plane and canceling current densities normal to the plane.

In this study, we investigated how large the effect of commonly used *in-vivo* probes is using the advanced EMI modeling framework. Our results are in line with these previous findings and we also show that the geometry, in terms of size and alignment of the probe, plays a very important role. We show that large silicon probes can be almost regarded as insulated planes when the neuron is aligned to them (potential increased by factor ~ 1.9) for large ranges of distances (Figure 5D). An interesting effect following the reduction of the amplitude factor with lateral shifts (Figure 6A) is that neurons not aligned with the probe will be recorded with a lower signal–to–noise ratio (SNR) due to the smaller amplitude increase, assuming that other sources of noise are invariant with respect to the probe location (such as electronic noise and biological noise from far neurons). This might bias neural recordings towards identifying neurons that are closer to the center of the probe, rather than the ones lying at the probes’ sides. However, this conclusion is speculative and might be affected by other factors, such as the distribution of neurons around the probe and their morphology (which contributes to the EAP). Therefore, ground truth information about the position of the recorded neurons and their reconstructed morphologies are needed for a quantitative evaluation of this phenomenon.

#### 4.2 Limitations and extensions

##### 4.2.1 Mesh improvements

The EMI model is, in principle, able to accurately represent the neuron and the neural probe. However, the accuracy of the model comes at the cost of computational resources. In order to be able to run simulations in a reasonable amount of time, the geometry of the neuron needed to be simplified considerably. First, we used a simple neuron in terms of a ball–and–stick with axon. This model is able to describe certain aspects of the neuronal dynamic [34], but it clearly cannot reach a level of detail of some more realistic morphologies, such as the reconstructed models made available by various initiatives [2, 1, 4, 5, 3]. We quantified the amplitude shift due to the probe in the extracellular domain (~ 1.9 on average for the MEA probes when neuron and probe are aligned), but this factor most likely also depend on the specific cell morphology that we used, and not only on the probe design and geometry. Therefore, we aim at extending the framework [49] for generating finite element meshes from publicly available realistic morphologies [5], allowing us to explore the *probe effect* for more complex morphologies.

Furthermore, we assumed *ideal* recording sites with an infinite input impedance which does not allow any current to flow in. In reality, recording electrodes have a high, but not infinite impedance that could be modeled by considering electrodes as an additional domain with very low conductance, even if it has been shown that for normal electrodes’ impedance the effect of conductive and equipotential recording sites is negligible [31].

##### 4.2.2 Computational costs

In Section 3.4 we showed that the EMI model is *much more* computationally demanding than conventional modeling using cable and volume conduction theory. For the simplest simulation performed in this study *(coarse 3* and *box size* 1), a system with 337515 unknowns was solved in about 40minutes. The Neuron simulations described in Section 2.4 took ~ 0.59 s to run, about 2400 times faster than the simplest EMI simulation performed here. However, because of our implementation and solution strategy for FEM, this factor should be considered as a rather pessimistic upper bound. In particular, the employed version of `FEniCS` (2017.2.0) does not allow for finite element spaces with components discretized on meshes with different topology. For example, the extra/intra-cellular potentials are defined on the entire Ω rather than Ω_*e*_ and Ω_*i*_ only, while the domain for the transmembrane potential *v* is Γ, but the space for *v* is setup on *all* facets of the mesh. For simplicity of implementation, the v unknowns on facets outside of Γ are forced to be zero by additional constraints and are not removed from the linear system. The LU solver thus solves also for the unphysical/extra unknowns and the memory footprint and solution times are naturally higher. The number of unphysical unknowns can be seen in Table 4 as a difference between total number of facets in the mesh and the number of facets on the surface of the neuron. For example, in the largest system considered here, avoiding the unphysical unknowns would reduce the system size by about 2 million.

In addition to assembling the linear system with only the physical unknowns, a potential speed up could be achieved by employing iterative solvers with suitable preconditioners. That is, fast PDE solvers for diffusion equations typically use around 1s per million degrees of freedom. As we here employ a H(div) formulation, we expect the solution to be computed in around 5 s per million degrees with multilevel methods. As shown in Table 4, 500 timesteps of solving systems with around one million degrees of freedom takes 82 600 s, which means 165 s per time steps. Hence, we may expect to speed up the solving procedure by around a factor 30 with better solvers. If further speed-up is required then finite element based reduced basis function method provides an attractive approach that should be addressed in future research.

##### 4.2.3 Finite elements methods are not alternatives to the conventional cable equation

The EMI framework, due to its computational requirements, is presently not an alternative to conventional modeling involving the cable equation (Equation 12) and the current summation formula (Equation 13). However, for specific applications, it can provide interesting insights. The hybrid solution combines the cable equation solution to finite element modeling, in practice solving the FEM problem only for the extracellular space and using the transmembrane currents computed by the cable equation as forcing functions [22, 46, 24, 20, 21]. However, the HS is also computationally expensive and it increases in complexity with longer simulation durations. Similar considerations can be made if Boundary Element Methods (BEM) [50] are employed instead of FEM ones, even though they are less computationally intense then the current FEM formulation. One possible drawback of BEM solvers is that they could not accommodate for anisotropic conductivity, while FEM solvers could in principle solve meshes with non-homogeneous conductivity between surfaces [51].

Another much faster option could be using method of images-based approaches [23, 24]. However, even correcting with a right factor smaller than 2, the MoI cannot account for the variability of peak ratios among the electrodes (Figure 8B). Therefore, we suggested here the probe correction (PC) method, which combines a one-time finite element simulation to model how each electrode of a specific probe affects the extracellular domain, and then uses the reciprocity principle to compute the potential on the recording sites arising from transmembrane currents. We showed that this method is able to reach the HS accuracy at a much smaller computational time (Table 4), which is also not strongly dependent on the simulation duration. Moreover, the time required to compute the probe specific mapping (*PC* (*map*)) and loading the FEM solutions in memory (*PC* (*load*)) could be further reduced by decreasing the mesh resolution. This possibility should be further investigated with a convergence analysis, similar to Section 3.2 for the EMI model.

#### 4.3 Significance of the *probe effect*

The effect of the recording device has not been fully taken into consideration in mathematical models of the extracellular field surrounding neurons. The *probe effect* needs to be considered when modeling silicon MEA, whose sizes are significantly larger than the recorded neurons. The assumption of an infinite and homogeneous medium is in fact largely violated when such *bulky* probes are in the extracellular space in the proximity of the cells. Although the tissue can be regarded as purely conductive and with a constant conductivity [52], these probes represent clear discontinuities in the extracellular conductivity, which strongly affect the measured potential due to their insulating properties. While the *probe effect* is large for MEAs, we found that it was negligible for microwire-type of probes, mainly for two reasons: first, the microwire is thinner and overall smaller than the MEA; second, the electric potential is sampled at the tip of the probe and in the entire semi-space below the microwire currents are free to flow without any obstacle.

When dealing with silicon MEAs, though, this effect could be crucial for certain applications that require to realistically describe recordings. For example, *Gold et al.* [25] used, in simulation, extracellular action potentials (EAP) to constrain conductances of neuronal models. Clearly, neglecting the *probe effect* would result in an incorrect parameterization of the models in this case.

Another example in which including this effect could be beneficial is when EAP are used to localize the somata position with respect to the probe. This is traditionally done by solving the *inverse problem:* a simple model, such as a monopolar current source [53, 54, 55], a dipolar-current source [53, 56, 57], line-source models [58, 59], or a ball–and–stick model [60], is moved around the extracellular space to minimize the error between the recorded potential and the one generated by the model. Ignoring the probe might result in larger localization errors.

Recently, we used simulated EAP on MEA as *ground truth* data, from which features were extracted to train machine-learning methods to localize neurons [26, 27] and recognize their cell type from EAPs [27]. When training such machine-learning models on simulated data and applying them to experimental data, neglecting the *probe effect* could confound the trained model and yield prediction errors.

Moreover, explaining experimental recordings on MEA without considering the probe might cause discrepancies between the modeling and experimental results hard to reconcile. On the other hand, in order to fully explain and validate our findings, an experiment with accurate co-location of extracellular recordings and cell position (and ideally morphology) is required. For example, an experimental setup in which a planar MEA is combined with two-photon calcium imaging [61] could provide an accurate estimate of the relative position between the neurons and the MEA.

In conclusion, we presented numerical evidence that suggests that the *probe effect,* especially when using Multi-Electrode silicon probes, affects the way we model extracellular neural activity and interpret experimental data and cannot be neglected for specific applications.

## Acknowledgments

A.P.B and K.H.J. are part of the Simula-UCSD-University of Oslo Research and PhD training (SUURPh) program, an international collaboration in computational biology and medicine funded by the Norwegian Ministry of Education and Research. T.N would like to acknowledge the European Union Horizon 2020 Research and Innovation Programme under Grant Agreement No. 720270 [Human Brain Project (HBP) SGA1].

## Footnotes

1 The reciprocity principle was originally derived for static charges and extended here to static currents.

